# β-alanine betaine and nAChRs in *Ascaris*

**DOI:** 10.64898/2026.06.30.735465

**Authors:** Paul D. E. Williams, David J. Borts, Dongjie Liu, Jacob Byerley-Duke, Brett VanVeller, Richard J. Martin

**Affiliations:** Department of Biomedical Sciences, Iowa State University, Ames IA 50011; Veterinary Diagnostic Laboratory, Iowa State University, Ames IA 50011; Department of Chemistry, Iowa State University, Ames IA 50011

**Keywords:** β-alanine betaine, nAChR, *Ascaris suum*, ligand, LC-MS/MS

## Abstract

Anthelmintic drugs are used to control soil-transmitted helminths that infect a third of the world’s human population. There is increasing concern about the development of resistance to anthelmintic drugs because of the limited number of compounds available and there is an unmet need for new resistance-busting drugs. Here we describe the presence of a previously unrecognized endogenous acetylcholine analogue, β-alanine betaine, which may serve as an endogenous ligand for an alternate subfamily of nicotinic receptors (DEG-3/DES-2) that could be developed as novel drug targets because their analogues are not present in their human or animal hosts. We collected peri-enteric fluid from female *Ascaris suum* (a model for the human parasite, *Ascaris lumbricoides*) and subjected it to chromatography and MS/MS to reveal signals consistent with acetylcholine, choline, and β*-*alanine betaine but we did not recover betaine. We injected betaine into female *Ascaris suum* which produced no effect. However, injection of β*-*alanine betaine, produced characteristic pretzel coiling and injection of levamisole produced a rod-like spastic paralysis. The differences between β*-*alanine betaine and levamisole suggested that they activate different nAChRs subfamilies. PCR showed that messages of the DEG-3 subfamily of nAChR channels, which are betaine targets and were present in the intestine and body wall of *A. suum*. Calcium signaling experiments showed that β*-*alanine betaine increased intracellular calcium of the intestine enterocytes and electrophysiology of the body muscle cells demonstrated that β*-*alanine betaine produced membrane potential depolarization. In N2 *elegans,* application of β*-*alanine betaine produced gradual inhibition of motility, which was reduced in *acr-20, acr-23, des-2, deg-3 and lgc-41* null-mutants. These observations suggest that, in addition to acetylcholine, β-alanine betaine - an anaerobic analog of betaine - may function as an endogenous ligand in anaerobic nematodes such as *A. suum*. An expanded repertoire of nicotinic acetylcholine receptor subfamilies in nematodes relative to mammals may reflect a corresponding need for diversification of cholinergic endogenous ligands in these organisms. This repertoire could allow their simpler neuronal system to perform more complex controls and be exploited for development of different and novel subfamily selective cholinergic anthelmintics.

**Author Summary:** There is increasing concern about the development of resistance to anthelmintic drugs because of the limited number of compounds available and there is an unmet need for new resistance-busting drugs. The cholinergic anthelmintics are one of the three major classes of anti-nematodal drugs that are used for control and treatment of soil-transmitted helminths. Each of these cholinergic anthelmintics (levamisole, pyrantel, derquantel, monepantel and oxantel) are selective for different nematode nicotinic acetylcholine receptors (nAChRs). The differences in selectivity could explain why resistance and species sensitivities varies across the different cholinergic anthelmintics. It is surprising how many nAChR genes are expressed in nematodes with more being present compared to humans. Why is this? Could it be that there are also more endogenous ligands other than acetylcholine allowing their simpler neuronal system to perform more complex control? We looked for additional analogues of acetylcholine in the body fluid of the large intestinal parasite of the pig *Ascaris suum* (a model for *Ascaris lumbricoides*) and identified the anaerobic cholinergic compound β-alanine betaine. We found evidence that suggests that β-alanine betaine may serve as an endogenous ligand for an alternate subfamily of nicotinic receptors (DEG-3/DES-2) that could be developed as novel drug targets because their receptor analogues are not present in human or animal hosts.

## INTRODUCTION

Soil transmitted helminths are common world-wide and affect a third of the human population Nematodes possess a large and diverse repertoire of nicotinic acetylcholine receptor (nAChR) subunit genes. In *C. elegans,* for example, there are ∼32 annotated subunit genes that have been classified into: the UNC-29 subfamily; the UNC-38 subfamily; the ACR-8 subfamily; the ACR-16 subfamily; and the DEG-3/DES-2 subfamily [1]. In addition, there are ∼25 orphan nAChR-like subunit genes that do not fall into these five core groups. In parasitic nematodes there are similar groupings of nAChRs and many subunit genes [2].

The cholinergic anthelmintics are one for the three major groups of therapeutic drugs used to treat soil-transmitted helminth infections. Each of the cholinergic anthelmintics (levamisole, pyrantel, derquantel, monepantel and oxantel) are selective for different nematode nAChRs that are each composed of different nAChR subunits [3, 4].

Of interest here is the DEG-3/DES-2 subfamily of nAChRs which in *C. elegans* includes ACR-5, ACR-17, ACR-18, ACR-20, ACR-23, ACR-24, DES-2, and DEG-3 subunits. Interest in the DEG-3/DES-2 family increased with the discovery of monepantel, an anthelmintic, which was found to modulate nAChRs with ACR-23 and ACR-20 subunits in *C. elegans* and MPTL-1 (Hc-ACR-22H) subunits in *Haemonchus contortus* [5–7]. Betaine and choline are endogenous nAChR ligands in *C. elegans* that function as agonists of ACR-20, ACR-23 and DEG-3/DES-2 nAChRs [6–8]. Apart from acetylcholine, it is not known if there are additional endogenous cholinergic ligands in parasitic nematodes.

We collected and analyzed constituents of *Ascaris suum* peri-enteric fluid looking for cholinergic ligands using HILIC–MS/MS and found signals for acetylcholine, choline, and β- alanine betaine but not betaine. Subsequent PCR experiments found multiple members of the DEG-3/DES-2 subfamily of nAChR subunits are present in the intestine and body wall of *Ascaris suum*. Intestine calcium signaling experiments and muscle bag electrophysiology found that β-alanine betaine increased enterocyte calcium and depolarized the body muscle. The paralytic effect of β-alanine betaine was not restricted to just parasitic nematodes as *C. elegans* became paralyzed when exposed to plates containing the compound, which was abated in nulls for different members of the DEG-3/DES-2 superfamily. These observations together suggest that in addition to acetylcholine, β*-*alanine betaine rather than betaine in the anaerobic nematode parasite may serve as an endogenous ligand and promote paralysis unlike the aerobic compound betaine. This large number of nAChR subunit genes found in nematodes and endogenous ligands compared to mammals may allow the simpler neuronal system of nematodes more complex control. The DEG-3/DES-2 superfamily of nAChRs is a distinctive group of ligand gated ion-channels that may be developed further as anthelmintic drug targets.

## METHODS

### Collection and maintenance of A. suum worms

Adult female *A. suum* worms were collected from the JBS Swift and Co. pork processing plant at Marshalltown, Iowa. Worms were maintained in *Ascaris* Ringers Solution (ARS: 13 mM NaCl, 9 mM CaCl_2_, 7 mM MgCl_2_, 12 mM C_4_H_11_NO_3_/ Tris, 99 mM NaC_2_H_3_O_2_, 19 mM KCl and 5 mM glucose pH 7.8) at 32 °C for 24 hrs. to allow for acclimatization before use in experiments. The solution was changed twice daily, and worms were used within three days of collection for experiments. All the worms were examined at the start of each day and excluded if they were damaged or became immotile.

### Injection of compounds into Ascaris suum

Healthy and motile *A. suum* were selected and placed in a dissection tray filled with warmed *Ascaris* Ringer’s solution. Worms were left to acclimatize in the tray for two minutes and were monitored for movement. All solutions were prepared in *Ascaris* Perienteric Fluid APF (23 mM NaCl, 110 mM NaAc, 24 mM KCl, 6 mM CaCl_2_, 5 mM MgCl_2_, 5 mM HEPES, 11 mM D-glucose, pH 7.6 using NaOH or Acetic acid). Worms were injected with either 500µL APF (negative control), 500µL 30 µM levamisole (positive control), 500µL 30 µM betaine or 500µL 30 µm β-alanine betaine. Solutions were injected into the worm above the collar using a 23-gauge needle attached to a 1ml Luer slip syringe. Worms were monitored for 2 minutes and were assessed for paralysis.

### A. suum cDNA synthesis and RT-PCR detection of DEG-3/DES-2 nAChR superfamily members

Intestine of *A. suum* was separated from the body wall which includes muscles, muscle bags and nerves, were homogenized separately in 1 ml of Trizol reagent using a mortar and pestle, followed by total RNA extraction according to the Trizol Reagent protocol (Life Technologies, USA). One microgram (1 µg) of total RNA from each tissue was used to generate cDNA by reverse transcription (RT) using SuperScript VILO^TM^ Master Mix (Life Technologies, USA) following the manufacturer’s protocol. PCR was conducted to detect the presence of *Asu-acr-20*, *Asu-acr-23*, *Asu-deg-3*, *Asu-des-2*, and *Asu-lgc-41* using primers targeting coding regions of each gene (Table 1)*. Asu-gapdh* was used as a reference gene. Negative controls included enzyme, water, and both forward and reverse primers for the target with no cDNA template. The cycling conditions for PCR were an initial denaturation for 2 min at 95 °C, followed by 35 cycles of 95 °C for 30 sec, 60 °C for 35 sec, 72 °C for 45 sec, and a final extension at 72 °C for 10 min using GoTaq^®^ G2 Hot Start Green Master Mix (Promega, USA). PCR products were separated on a 2% Agarose gel containing SYBR^®^ Safe DNA Gel Stain, followed by visualization and images were captured using an Azure 600 imaging system set to SYBR Safe (Excitation 472nm, Emission 595nm). Uncropped and unedited raw images are shown in Supplemental Fig. S1.

**Table 1.**
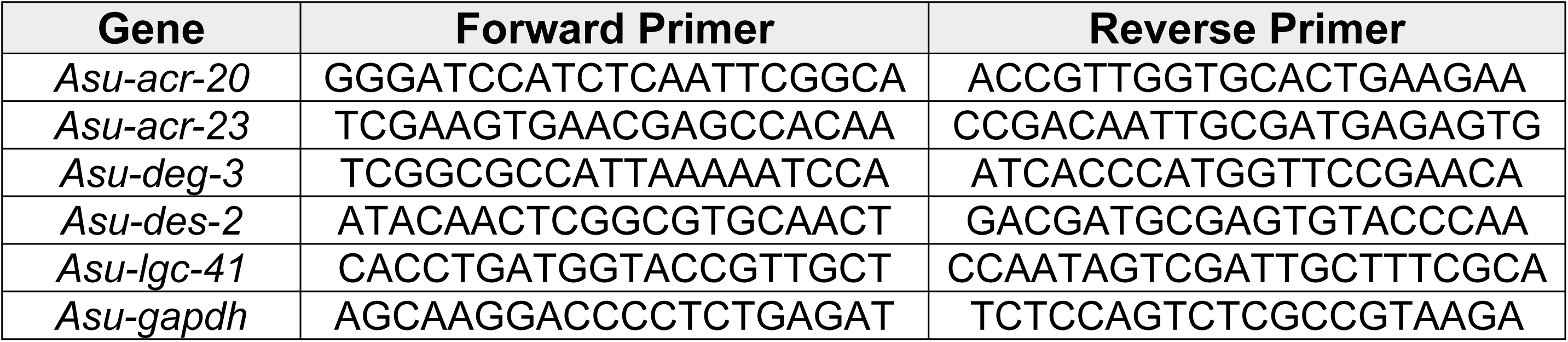
Primer sequences used for RT-PCR. Forward and reverse primer sequences used for the detection of *acr-20, acr-23, deg-3, des-2, lgc-41* and the reference gene *gapdh* in *Ascaris suum*.

### C. elegans strains and maintenance

*C. elegans s*trains were maintained on NGM agar plates seeded with *E. coli* OP50 bacteria per standard protocols. Strains used were: N2, RB2119 *acr-23(ok2804) V*, VC1598 *acr-20(ok1849)/mT1 II; +/mT1 [dpy-10(e128)] III*, TU1803 *deg-3(u662); des-2(u695) V* and PS8729 *lgc-41(sy1494) X*. All strains were acquired from the Caenorhabditis Genetics Center (University of Minnesota, Minneapolis, MN, USA).

### Ascaris suum muscle flaps for electrophysiology

1 cm muscle tissue flaps were prepared by dissecting the anterior part of the worm, 2-3 cm caudal to the head. A body muscle flap preparation was then pinned onto a Sylgard^™^-lined double jacketed bath chamber maintained at 35°C by inner circulation of warm water (Fisher scientific Isotemp3016H, Pittsburgh, PA, USA). The intestine was removed to expose the muscle cells [9]. The preparation was continuously perfused, unless otherwise stated, with APF. The incoming perfusate was pre-warmed to 35°C with an in-line heating system (SH 27B Warner instruments, Hamden, CT, USA) before it was perfused at 3.5-4 mL/min through a 20-gauge (1.5″ long) needle placed directly over the muscle bag recorded from. The experimental compounds were dissolved in APF as described in the results. 30 µM β-alanine betaine, 100 µM β-alanine betaine, 300 µM β-alanine betaine and 1 mM β-alanine betaine were applied for a period of 10 seconds by means of a tube that rapidly replaced all the bathing solution covering the *Ascaris* muscle cells. Flanking applications of 10 µM acetylcholine (ACh) were applied at the start and end of each recording for 10 secs to determine viability.

A single micropipette was used to record the membrane potential and to examine the electrophysiological effects in the *A. suum* muscle bag region. Borosilicate capillary glass (Harvard Apparatus, Holliston, MA, USA, ID-0.86mm, OD- 1.5mm) micropipettes were pulled on a Flaming Brown Micropipette puller (Sutter Instrument Co., Novato, CA, USA) and filled with 3M potassium acetate (resistance 20-30 MΩ). The recordings were obtained by impaling the bag region of *A. suum* muscle with both the micropipettes. All experiments were performed using an Axoclamp 2A amplifier, a 1320A Digidata interface and Clampex 9 software (Molecular Devices, Sunnyvale, CA).

### C. elegans paralysis assays

Motility responses to β-alanine betaine were assayed on NGM plates that were prepared two hours before the experiment. 50 mM β-alanine betaine was added to molten agar (∼55°C). Once solidified fresh *E. coli* OP50 was added to the plate and were left to air dry. 20-40 L4 animals were picked the night before the assay onto fresh OP50 seeded NGM plates and left to develop to young adults overnight at room temperature. Animals were transferred from the stock plate to an intermediate plate for one minute to allow animals to separate and be assessed for motility. A total of ten suitable animals were then transferred to β-alanine betaine assay plates and motility was measured. Worms that either appeared damaged, had reduced movement or no movement were removed from the assay plate and replaced with viable *C. elegans*. Animals were allowed to free roam on the bacterial lawn and were assessed after 10, 20, 30, 45, 60, 90 and 120 minutes. Animals were scored based on movement by lightly tapping the ‘nose’ of the worm with a hair. If the worm failed to respond to the hair, they were considered paralyzed.

### Preparation and loading Fluo-3AM

A 2 cm section of the intestine was removed from the body piece using fine forceps and cut open. The intestinal flap was placed onto a coverslip (24 x 50 mm) and pinned using a slice anchor (26 x 1mm x 1.5mm grid, Warner Instruments, Hamden, CT), immersed in 1 mM CaCl_2_ APF (23 mM NaCl, 110 mM NaAc, 24 mM KCl, 1 mM CaCl_2_, 5 mM MgCl_2_, 5 mM HEPES, 11 mM D-glucose; pH 7.6) in a laminar flow chamber (Warner RC26G, Warner Instruments, Hamden, CT). Fluo-3AM (Sigma-Aldrich, St. Louis, MO, USA) loading was achieved by incubating the tissue in Ca^2+^-free (>100 µM) APF containing 5 µM Fluo-3AM and 0.05% Pluronic F-127 (10% v/v in water; Invitrogen, Thermo Fisher Scientific, Waltham, MA, USA) for 60 minutes with the recording chamber connected to a Dual Automatic Temperature Controller (Warner Instruments, Holliston, MA, USA) maintained at 36-37°C. After incubation, the Fluo-3AM solution was discarded, and the sample was incubated in 1 mM CaCl_2_ APF for an additional 20 minutes at 36-37°C to promote calcium loading. Fluo-3AM loading was confirmed under blue light using a blue/green filter box (EF-4 AT EGFP/FITC/CY2/ALEXA FLUOR Band Pass; Excitation: 465-495nm, Emission: 525-545nm, Dichroic Mirror) (Chroma Technology, Bellows Falls, VT, USA) and visualized under pseudo-color settings using MetaFluor 7.10.2 (Figs. 2A & B). Any tissue that did not show Fluo-3 fluorescence was discarded.

### Measurement of Ca^2+^ fluorescence

All recordings were performed on a Nikon Eclipse TE3000 microscope fitted with a 20X/0.45 Nikon PlanFluor objective, using a Photometrics Retiga R1 Camera. Light control was achieved using a Lambda 10-2 with a shutter controller. Fluorescence was achieved using a Lambda LS Xenon bulb lightbox (Sutter Instruments, Novato, CA, USA) which delivered light via a fiber optic cable to the microscope which passed through a blue/green filter box. Fluorescent light emission was controlled by using the shutter. Minimal illumination exposure was used to prevent photobleaching.

During recordings tissues were continuously perfused with 1 mM CaCl_2_ APF buffer solutions. Intestinal preparations were exposed to 1 mM β-alanine betaine for five minutes. Application of 10 mM CaCl_2,_ which was used as a positive control to determine tissues viability, was applied 5 minutes after the β-alanine betaine signal returned to baseline to account for any delayed or latent responses to occur. If a preparation failed to respond to the 10 mM CaCl_2_ control signal it was discarded from the data pool regardless of previous signals to β-alanine betaine. Every experiment reported in this study had a 100% response to 10 mM CaCl_2_. A change in fluorescence ≥5% was classified as a positive response to a compound as the average spontaneous change in cells continuously perfused with APF buffer was ∼2.5% [10]. Stocks and working concentrations of all compounds were made in 1 mM CaCl_2_ APF buffer solution.

All solutions were delivered to the chamber under gravity feed through solenoid valves controlled using a VC-6 six-channel Valve Controller through an inline heater set at 37°C (Warner Instruments, Holliston, MA, USA), at a rate of 1.5mL/min. At the start of all experiments, intestinal preparations were left under blue light for a minimum of 3 minute to promote settling and equilibration of the fluorescent signal and to monitor any spontaneous calcium signals.

All calcium signal recordings were acquired and analyzed using MetaFluor 7.10.2 with exposure settings at 250 ms with 2x binning. Calcium signals from each intestine were collected from 50 square 50 µm x 50 µm areas across the intestine covering a total area of 125,000 µM^2^ that included 800-1000 individual enterocytes. The average fluorescence amplitude was calculated for each intestinal exposure of all 50 regions. Maximal percent calcium signal amplitudes (ΔF) were calculated using the equation F1-F0/F0 x 100, where F1 is the fluorescent value and F0 is the baseline value. All F0 values were taken at the time point each compound was applied for every cell analyzed. All representative response traces are presented as the average percent change in calcium fluorescence with the standard error of the mean (±SEM) of all cells from a single recording. Traces were generated by converting the calcium signal profiles of all the regions from a single recording to percentages using the previously described ΔF/F0 equation, with the time of stimulus application being F0 (0% for all cells) and 100% being the peak value for each cell. The mean percentage change in fluorescence and the SEM was calculated for each time point during the application of each compound. All baseline values were taken at the time point each compound was applied for every cell. Traces are represented as mean ±SEM. The number of intestinal tissues recorded (n), along with the total number regional responses are provided in the figure legends.

### Determination of choline analogues by HILIC–MS/MS

2.5 ml *Ascaris suum* peri-enteric fluid samples were collected from freshly collected worms and 20 µM neostigmine was added as a choline esterase inhibitor. Choline analogues in *Ascaris suum* peri-enteric fluid were identified using hydrophilic interaction liquid chromatography coupled to full mass range (HILIC-MS) and tandem mass spectrometry (HILIC–MS/MS).

### Sample Preparation

Fluid samples were maintained on ice and mixed thoroughly. Proteins were precipitated by adding three volumes of ice-cold acetonitrile. Samples were vortexed for 30 s and incubated on ice for 10 min. The mixture was centrifuged at 15,000 × g for 10 min at 4 °C. The supernatant was transferred to autosampler vials and diluted with acetonitrile to achieve 80–90% organic solvent prior to injection.

### HILIC Chromatography

Chromatographic separation was performed using a Thermo Fisher Scientific Vanquish Flex UHPLC system with a HILIC column (Agilent Infinity Lab Poroshell 120 HILIC-Z; 2.1 x 100 mm, 2.7 µm particle size) maintained at 30°C. The mobile phases consisted of: Mobile phase A: 95:5 Water/Acetonitrile with 10 mM Ammonium Acetate and Mobile phase B: 95:5 Acetonitrile/Water with 10 mM Ammonium Acetate. The gradient started at 90% B, ramped linearly to 50% B over 5 minutes, then from 50% B to 40% B over 1 minute, held for 2 minutes at 40% B, then returned to initial conditions over 1 minute, and equilibrated for 6 min. The flow rate was 0.3 mL/min, and the injection volume was 10 µL.

### Mass Spectrometry

Detection was performed using a Thermo Fisher Scientific Exploris 120 quadrupole-orbitrap mass spectrometer equipped with an electrospray ionization (ESI) source operated in positive ion mode. The following source parameters were used: capillary voltage 3.5 kV, Sheath Gas = 50 (Arb), Aux gas = 10 (Arb), Sweep gas = 1 (Arb), Ion Transfer Tube Temp = 325 °C, Vaporizer Temp = 350 °C.

Additional mass spectrometer settings include: full mass range resolution = 60,000, MS/MS resolution = 15,000, full mass range = 50 – 500 *m/z*, MS/MS mass range was set to “auto”, Automatic Gain Control (AGC) was set to ‘standard’ for both full mass range and MS/MS, and maximum ion time was set to ‘auto’ for both full mass range and MS/MS. MS/MS data was acquired in t-MS2 (targeted MS2) mode.

### Statistical Analysis

Statistical analysis of all data was done using GraphPad Prism 9.0 (GraphPad Software, Inc., La Jolla, CA, USA). To ensure reproducibility, we repeated our experiments: the numbers of animals, number of preparations, the concentrations, and durations of applications are provided in the legends of the figures. Analysis of changes in depolarization were done using paired or unpaired student *t*-tests with *P < 0.05* being considered significant. All data are presented as mean ± SEM for each treatment.

### Chemicals

Source of chemicals: β-alanine betaine used in all experiments was produced by ISU and characterization matched previous reports [11], except for the neat chemical standard which was purchased from Smolecule. Betaine was supplied by Spectrum Chemicals and levamisole was provided by MP biomedicals. Fisher Scientific supplied all other chemicals.

## RESULTS

There are a surprising number of nAChR subunit genes in nematodes. There are ∼32 annotated subunit genes in *C. elegans* that have been divided into five subfamilies: 1) the UNC-29 subfamily, 2) the UNC-38 subfamily, 3) the ACR-8 subfamily, 4) the ACR-16 subfamily, and 5) the DEG-3/DES-2 subfamily [1]. The DEG-3/DES-2 subfamily group are interesting because some of their nAChRs have been activated by the anthelmintic monepantel, betaine and/or choline [5]. We wondered if there could be additional unrecognized endogenous cholinergic ligands present in nematodes that were selective for these different nAChRs. We describe here observations that suggest that β*-*alanine betaine (Fig. 1) may be an additional endogenous cholinergic ligand in *Ascaris suum*.

**Fig. 1:**
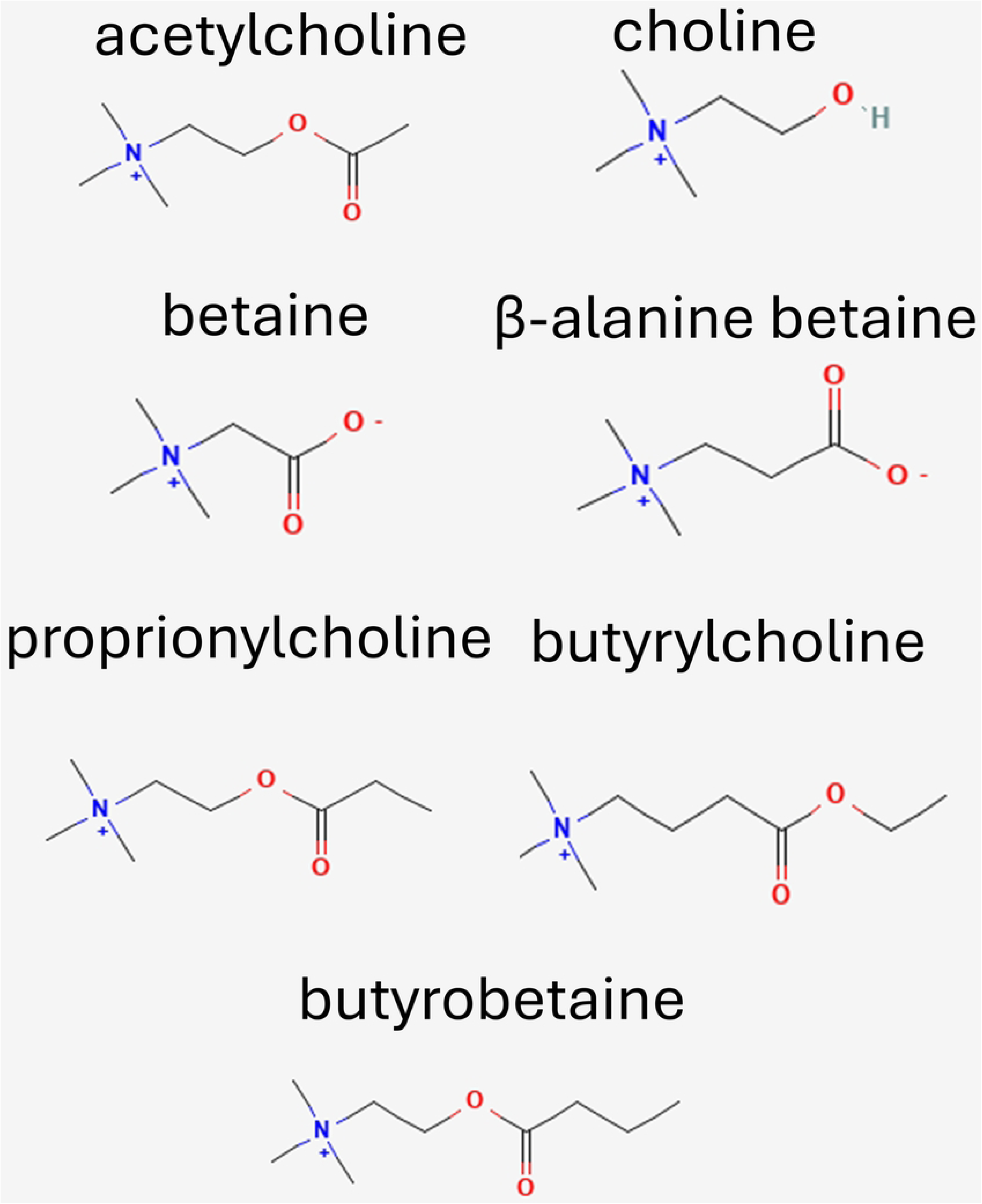
Chemical structures of acetylcholine, choline, betaine, β-alanine betaine, proprionylcholine, butyrylcholine and butyrobetaine.

### Collection of peri-enteric fluid and identification of β-alanine betaine

The pseudocoelomic fluid of *A. suum* is a protein rich, metabolically active fluid that distributes metabolites and nutrients along the length of the nematode. This fluid is found between the intestine and cuticle of the worm and bathes the muscle and nerves. Signaling molecules including cholinergic transmitters like acetylcholine can be found in this fluid. To determine if cholinergic transmitters are present in the pseudocoelomic fluid, we dissected adult female *A. suum* (N = 4) and drained the fluid from the parasite. We added 20 µM neostigmine to inhibit esterases that could degrade any cholinergic compounds in the fluid.

The fluid was analyzed using Hydrophilic Interaction Liquid Chromatography (HILIC) Liquid Chromatography-Mass Spectrometry (LC-MS). We observed signals consistent with acetylcholine and choline (not shown) but did not observe evidence of betaine. Acetylcholine, synthesized from choline, is the principal excitatory neurotransmitter in nematodes and is extensively characterized through biochemical and electrophysiological studies. Choline has been identified previously in the nematodes *C. elegans* [12]; *Necator Americanus* and *Nippostrongylus braziliensis* [13]. Betaine has been identified previously in *C. elegans* [14]; *H. contortus* [15]; *Necator Americanus;* and *Nippostrongylus braziliensis* [13, 16]. But we did not find evidence of betaine in *A. suum.* Interestingly, we identified an unknown compound in the pseudocoelomic fluid Fig. 2A with a retention time of 3.39 min. After further analysis, the compound was hypothesized to be β-alanine betaine, Fig.1.

**Fig. 2:**
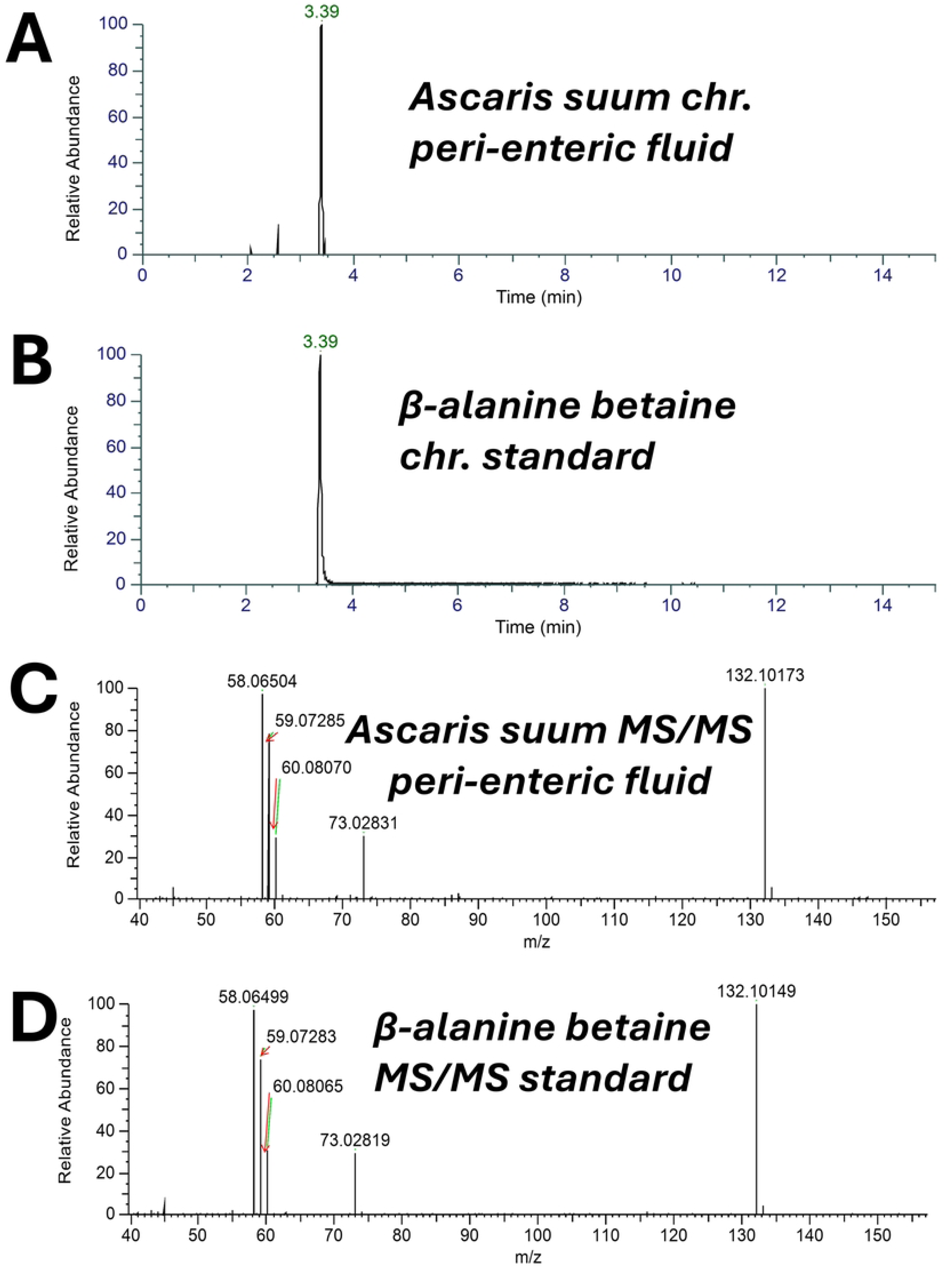
Chromatograms and HILIC–MS/MS showing signals for β-alanine betaine. A: Chromatogram from *A. suum* pseudocoelomic fluid for suspected β-alanine betaine. B: Chromatogram for β-alanine betaine neat chemical standard. C: MS/MS spectrum for suspected β-alanine betaine from *A. suum* pseudocoelomic fluid. D: MS/MS spectrum for β-alanine betaine neat chemical standard.

To verify that the unknown compound was β*-*alanine betaine, a standard was purchased from Smolecule, and a second round of HILIC LC-MS was performed. The β-alanine betaine standard matched the profile of the presumed β-alanine betaine from the pseudocoelomic fluid with an identical retention time of 3.39 min, Fig. 2B, and Tandem Mass Spectrometry peaks at 58.06, 59.07, 60.08, 73.03, and 132.10 *m/z*, Figs. 2C & D. Signals consistent with β*-*alanine betaine were observed in all 4 samples. We did not detect any other signals that were consistent with other choline or betaine derivatives including: propionylcholine, betaine, butyrylcholine, or butyrobetaine in any of the 4 samples.

### Pretzel effect of injected β-alanine betaine

With the detection of β*-*alanine betaine in the peri-enteric fluid of *A. suum,* we wanted to determine if this compound played a role in modulating motility, like other cholinergic compounds. We injected either 500 µL of APF salt solution (negative control: N=4), 500 µL 30 µM levamisole (positive control: N=4), 500 µL 30 µM betaine: N=4, or 500 µL 30 µM β*-*alanine betaine: N=4, into separate free moving adult female *A. suum* using a fine needle, just in front of the gonopore and compared the effects on motility. The APF injection did not affect the continuous slow movement of any of the worms or affect their posture, Fig. 3A. The injection of levamisole rapidly inhibited movement and all worms straightened into rod-like structures, Fig. 3B. The injection of betaine, like that of the APF injection did not affect the slow movement or posture of the worms, Fig. 3C. However, the effect of β*-*alanine betaine was different to the other compounds as after injection, as all worms slowly became coiled and formed a ‘pretzel’ like structure, Fig. 3D. These results demonstrate that β*-*alanine betaine promotes a unique paralytic phenotype in *A. suum* by potentially acting on cholinergic receptors.

**Fig. 3:**
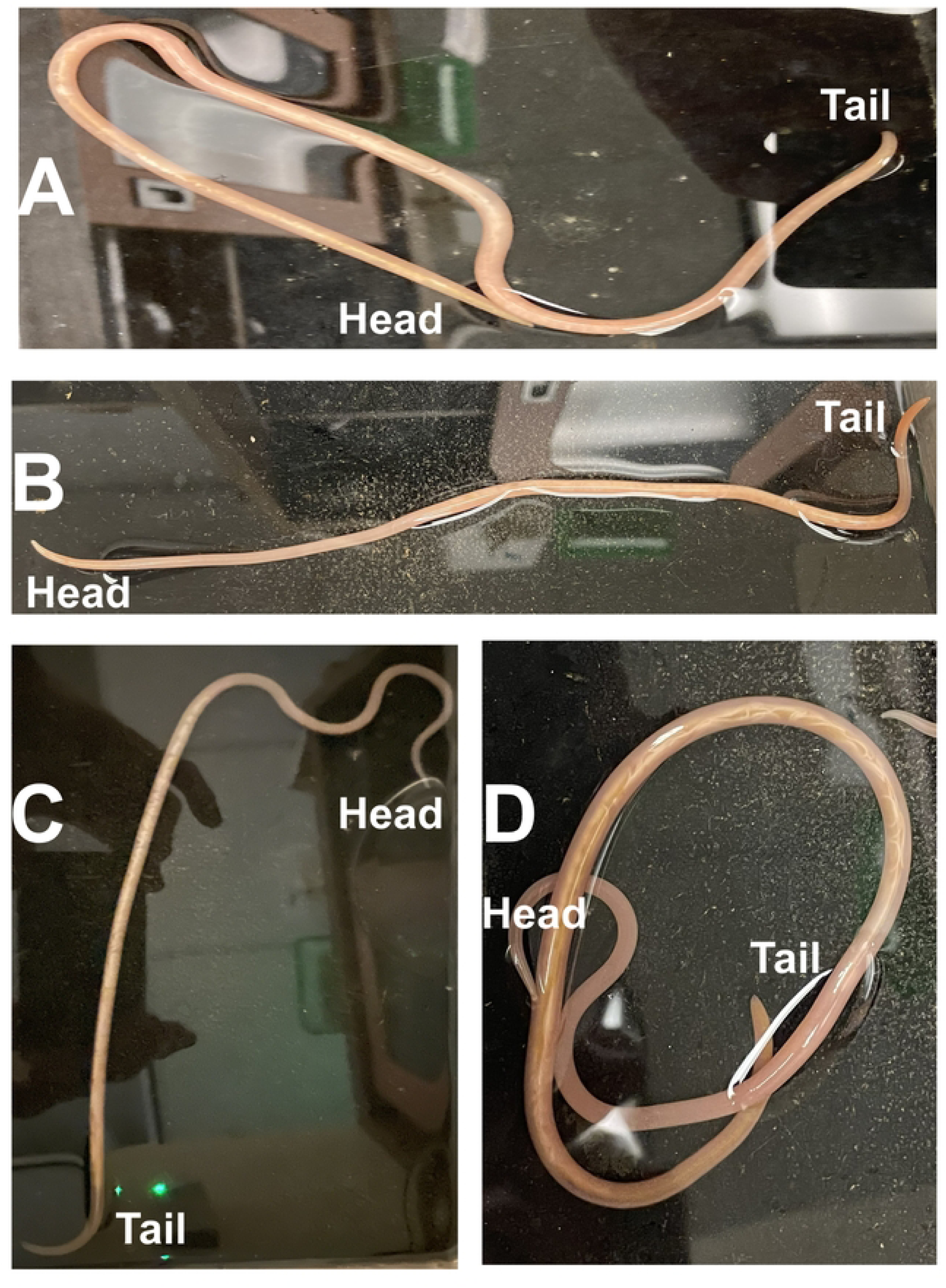
Injection of β-alanine betaine promotes “pretzel” paralysis phenotype. A: Adult female *A. suum* injected with 0.5mL APF, the worm continued to move with no clear effect, n = 4. B: Adult female *A. suum* injected with 0.5mL 30 µm levamisole: note the straight rod paralyzed appearance, n = 4. C: Adult female *A. suum* injected with 0.5mL 30 µm betaine: the worm continued to move with not striking effects, n = 4. D: Adult female *A. suum* injected with 0.5mL 30 µm β-alanine betaine. Note the coiling (pretzel) posture, n = 4.

### DEG-3/DES-2 subfamily subunits, including ACR-23, are present in Ascaris intestine and body wall

The different effects of injected levamisole and injected β*-*alanine betaine on *A. suum* paralysis suggests the two compounds act at different sites. Levamisole activates nAChRs which are composed of UNC-29, UNC-63, ACR-8, UNC-38 subunits which are not part of the DEG-3/DES-2 subfamily [17]. In *C. elegans,* betaine has been established to be an agonist of either homomeric nAChRs composed of ACR-23 or ACR-20 [18], DEG-3/DES-2 heteromeric receptors [8], or the betaine-gated chloride channel LGC-41 [19], suggesting that the β*-*alanine betaine could act on similar nAChR subunits in *A. suum*.

In *C. elegans*, ACR-23 has been found to be expressed in different tissues, including body wall muscles. To determine if *acr-23* is present in *A. suum,* we performed a blast search with *Cel-acr-23* against the *A. suum* genome and identified the gene AgR001_g108, as the *acr-23* orthologue with 53.6% similarity in the protein sequence to Cel-ACR-23. We generated primers targeting *Asu-acr-23* and screened for the presence in paired cDNA pools of intestine and body wall (the body wall includes the muscle and nerve cords). We found that *Asu-acr-23* is present in both cDNA pools, Fig. 4 (and S1) suggesting global expression of the receptor. To ensure that we did successfully target *Asu-acr-23* we performed full length sequencing of the AgR001_g108 from our cDNA pools using Sanger sequencing and compared our results to the database and obtained 99.8% sequence similarity.

**Fig. 4:**
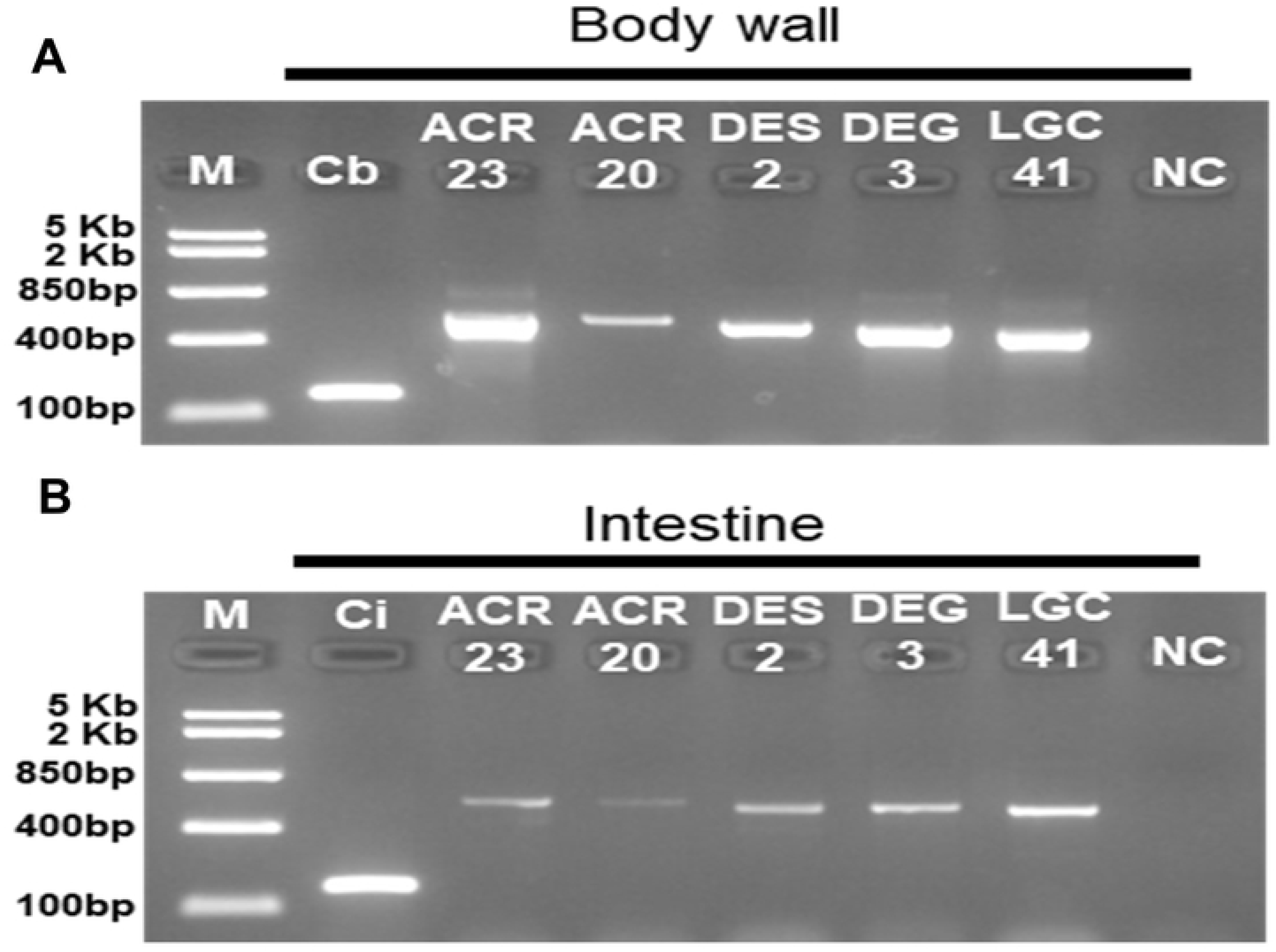
RT-PCR analysis of Asu-acr-23, acr-20, des-2, deg-3 and lgc-41. A: paired body wall B: intestine from female *A. suum*. Each lane represents one of the genes for the intestine or body wall of an individual worm. *Asu-gapdh* from the intestine or body wall was used as a positive control. Neg. = negative control, no cDNA template present. M = FastRuler Middle Range DNA Ladder (ThermoFisher Scientific).

With the global detection of *Asu-acr-23,* we identified and screened for the presence of *A. suum* orthologues for the other *C. elegans* betaine sensitive channels including: ACR-20, DES-2 and DEG-3 subunits and the inhibitory betaine activated ion-channel LGC-41 as possible β*-*alanine betaine targets (Table 2). We detected the presence of all four subunits in both the intestinal and body wall cDNA pools of *A. suum* Fig. 4 (and S1), with *Asu-acr-23* having the brightest band in the body wall. These results suggest that targets composed from these genes may mediate the effects of *β-*alanine betaine in *A. suum*.

**Table 2:** Genes and accession numbers for members of DEG-3/DES-2 nAChR superfamily in A. suum. Note *Asu-deg-3* and *Asu-des-2* had strong similarity with AgR001_g107 but had different predicted isoforms.

| Gene | Gene Identifier | Accession number |
| --- | --- | --- |
| *Asu-acr-20* | AgR001_g020 | AEUI03000002.1 |
| *Asu-acr-23* | AgR001_g108 | AEUI03000002.1 |
| *Asu-deg-3* | AgR001_g107_t06 | AEUI03000002.1 |
| *Asu-des-2* | AgR001_g107_t03 | AEUI03000002.1 |
| *Asu-lgc-41* | AgR002_g403_t03 | AEUI03000004.1 |

### Intestine and muscle cells responses to β-alanine betaine

With the identification of potential receptors for β-alanine betaine, we wanted to determine if application to either the intestine or muscle elicited a response. We first tested the effects of β*-*alanine betaine on the intestine of *A. suum* by measuring the Ca^2+^ signal. We found that 1 mM concentrations of β*-*alanine betaine induced large and prolonged increases in enterocyte intracellular Ca^2+^, with an average amplitude of 31% ± 2.09% (N = 3), Figs. 5A & B. Interestingly the increase in the Ca^2+^ signal did not occur immediately after applying β*-*alanine betaine but instead took a couple of minutes before the Ca^2+^ started to increase Fig. 5A. The Ca^2+^ signal started to slowly decline after the β*-*alanine betaine was washed out before returning to near baseline levels. We observed that 79% ± 8.1% of the intestinal regions tested were stimulated by the β*-*alanine betaine Fig. 5C, suggesting that the receptors were not localized to specific regions of the intestine. To test for viability, 10mM CaCl_2_ was applied and we observed robust increases in Ca^2+^ (54% ± 1.8%) in 100% of regions tested Figs. 5B & C.

**Fig. 5:**
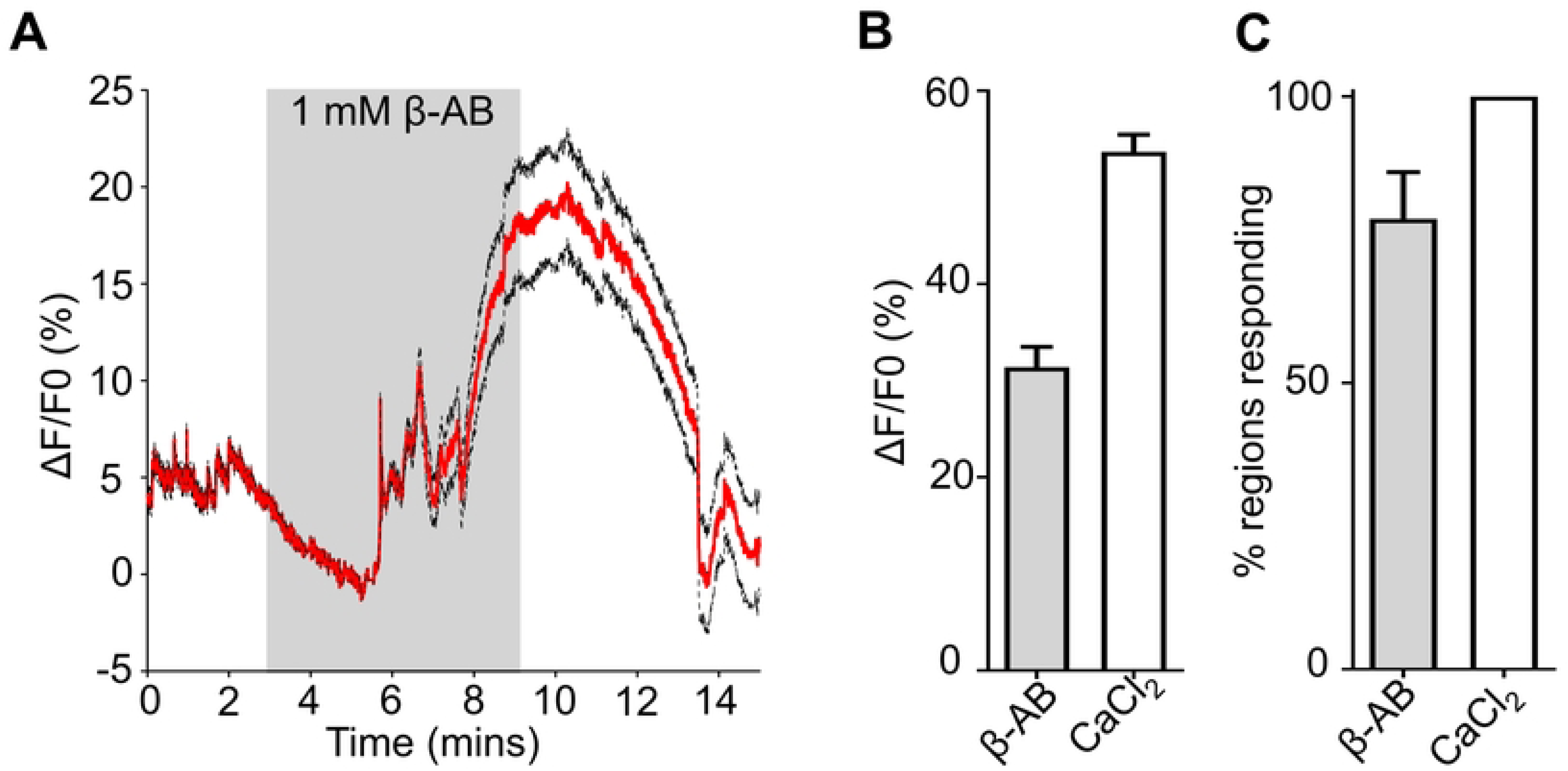
β-alanine betaine stimulates robust increases in Ca^2+^ in *A. suum* intestines. A: Representative response to 1mM β-alanine betaine. Red lines indicate average fluorescence; black dotted lines represent ±SEM. Grey Box indicates stimulus application. B: Total average maximal amplitudes of Ca^2+^ fluorescence to 1mM β-alanine betaine (grey bar) and 10mM CaCl_2_ (white bar). C: Mean percent of regions showing a Ca^2+^ response to 1mM β-alanine betaine (β-alanine betaine: grey bar) and 10mM CaCl_2_ (white bar). *n* = 3 intestines from 3 individual female *A. suum* with 150 total regions. β-alanine betaine 119/150 regions responding, CaCl_2_ = 150/150 regions responding. All values represented as means ± SEM.

We also measured the effects of β*-*alanine betaine on the membrane potential of *A. suum* body muscle cells. We applied 10 sec applications of 30 µM, 100 µM, 300 µM and 1 mM β*-*alanine betaine to the body muscle cells and compared the effects with flanking 10 µM acetylcholine applications, Fig.6. In each of the muscle cell preparations, the depolarization in response to β*-*alanine betaine was slow and significantly weaker but demonstrated a concentration dependent effect, with 30 µM β*-*alanine betaine having an average ΔmV of 0.06 mV ± 0.03, 100 µM having a ΔmV of 0.4 ± 0.2, 0.3 mM having an average ΔmV of 1.2 mV ± 0.3 and 1 mM having an average of 2.2 mV ± 0.4 Fig. 6A & B. We observed no significant difference between the flanking 10 µM acetylcholine applications (6.9 mV ± 1.1 & 7.1 mV ± 0.9) Fig. 6B. We also observed a concentration dependent effect on the rate of spontaneous depolarizations (spikes) that took more than a minute before reaching its full effect with 1 mM β*-*alanine betaine having the highest frequency 79.3 (± 16.4) Fig. 6A & C. Again, we observed no significant difference between the flanking 10 µM acetylcholine applications (25.5 ± 5.1 & 25.75 ± 4.1) Fig. 6C. Together these results demonstrate that β*-*alanine betaine has effects on *A. suum* muscle depolarization and intestinal calcium signaling suggesting a novel functional cholinergic ligand in an anerobic parasite.

**Fig. 6.**
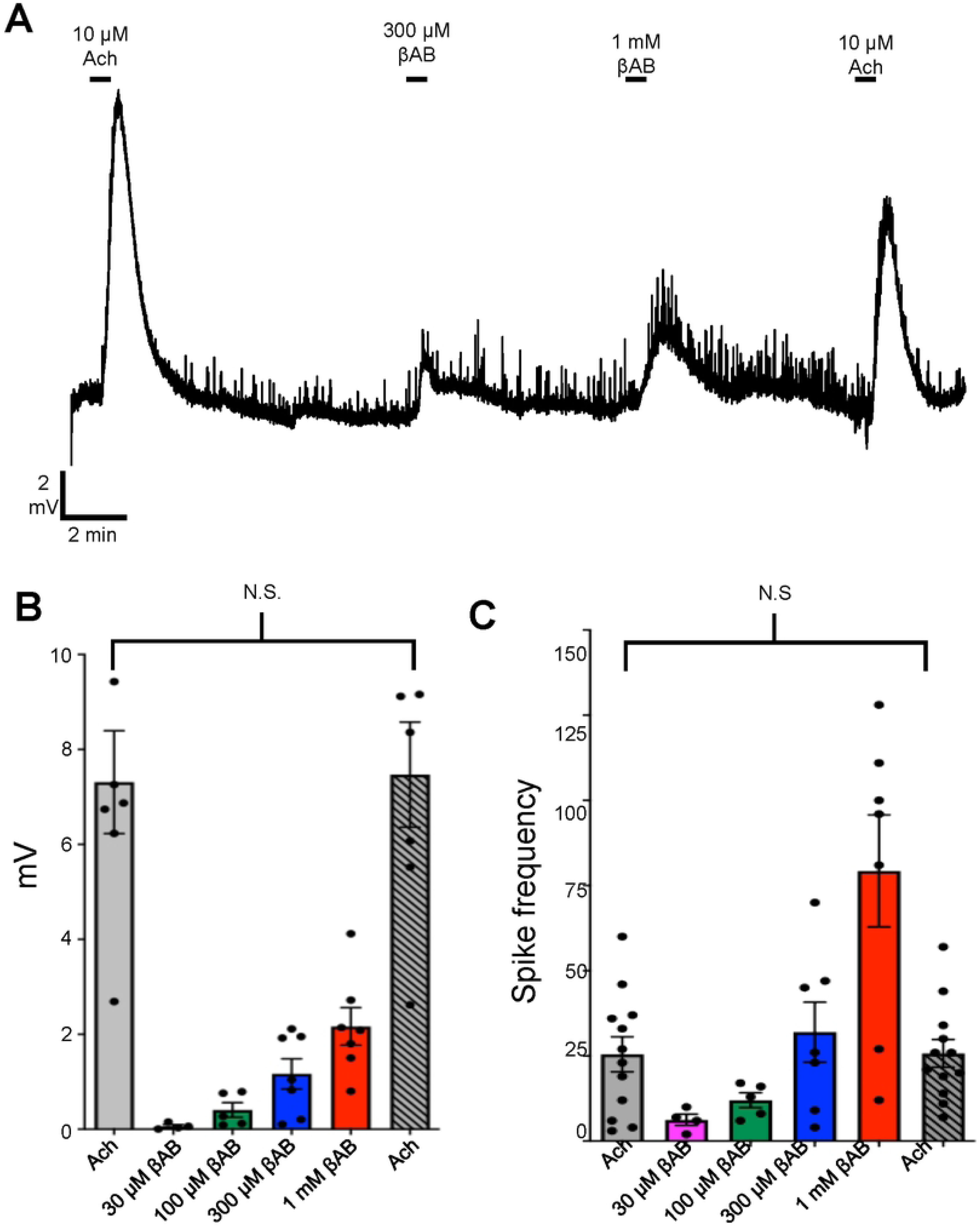
β-alanine betaine stimulates small depolarizations on the muscle bags of *A. suum*. A: An intracellular membrane potential recording from the muscle bag-region of a female *A. suum* worm to higher concentrations of β-alanine betaine. Sample was first exposed to an application of 10 µM acetylcholine by rapid perfusion for 10sec followed by an application of 300 µM β-alanine betaine for 10 sec, 1 mM β-alanine betaine for 10 sec and finally to a second application of 10 µM acetylcholine for 10 sec. B: Histograms of the mean +/- S.E.M. peak depolarization responses to the first 10 µM acetylcholine application (grey bar), 30 µM β-alanine betaine, 100 µM β-alanine betaine (green bar), 300 µM β-alanine betaine (blue bar), 1 mM β-alanine betaine (red bar), and the second application of 10 µM acetylcholine (grey bar; hashed). N.S. not significantly different to 1^st^ 10 µM acetylcholine (1^st^ 10 µM acetylcholine vs 2^nd^ acetylcholine, *P* = 0.799*, t* = 0.2613, *df* = 11, paired *t*-test). C: Histograms of the mean +/- S.E.M. number of spikes produced during the application of first 10 µM acetylcholine application (grey bar), 30 µM β-alanine betaine, 100 µM β-alanine betaine (green bar), 300 µM β-alanine betaine (blue bar), 1 mM β-alanine betaine (red bar), and the second application of 10 µM acetylcholine (grey bar; hashed). N.S. not significantly different to 1^st^ 10 µM acetylcholine (1^st^ 10 µM acetylcholine vs 2^nd^ acetylcholine, *P* = 0.931*, t* = 0.088, *df* = 11, paired *t*-test). ACh recordings *n =* 12 individual muscles from 12 individual female *A. suum* females. 300 µM & 1 mM β-alanine betaine *n* = 7 total recordings from 7 individual *A. suum* females. 100 µM β-alanine betaine *n* = 5 total recordings from 5 individual *A. suum* females. 30 µM β-alanine betaine *n* = 4 total recordings from 4 individual *A. suum* females. All values represented as means ± SEM.

### Effects of β-alanine betaine and DEG-3 C. elegans subfamily null mutants

With the evidence that β*-*alanine betaine affects the movement and produces a pretzel shaped *A. suum* after injection, we utilized the free-living nematode *C. elegan*s to determine if an effect of β*-*alanine betaine was nematode species specific. Following previous techniques with betaine and using established techniques with C*. elegans* with high concentrations that are required to cross their thick cuticle and to overcome bacterial metabolism [20, 21], we incubated wild-type N2 *C. elegans* animals on *E. coli* OP50 seeded NGM plates containing 50 mM β*-*alanine betaine. We observed that the motility of *C. elegans* decreased progressively over 2 hours with 77% of the control N2 *C. elegans* paralyzed by the end of the experiment Fig. 7A: orange line. The N2’s lost their stereotypical sinusoidal wave phenotype Fig. 7B. With ACR-23 being responsible for betaine mediated signaling in *C. elegans*, we hypothesized that β*-*alanine betaine would act on similar nAChRs. We exposed *acr-23(ok2804)* nulls to 50 mM β*-*alanine betaine and observed reduced paralysis compared to N2, with only 19% of animals being paralyzed after 2 hours Fig. 7A: red line. Unlike N2, *acr-23* nulls still maintained the sinusoidal waveform Fig. 7C. We repeated the experiments using 50 mM betaine and saw no evidence of larval arrest or animal paralysis like previous publications [6,19]. These results suggest that β-alanine betaine paralyzes *C. elegans* with effects mediated via ACR-23 receptors.

**Fig. 7.**
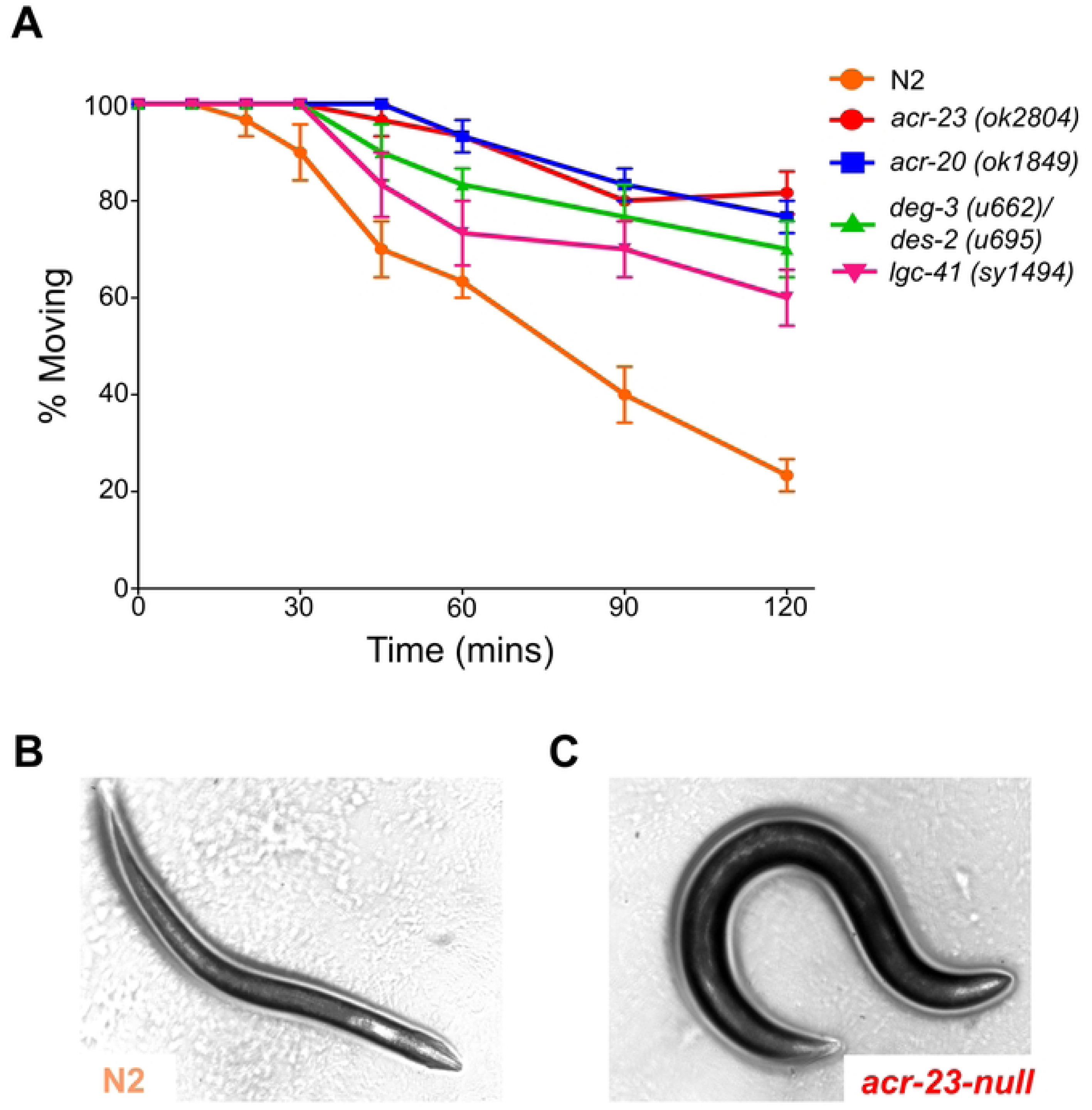
β-alanine betaine paralyzes *C. elegans* through DEG-3/DES-2 subfamily members. A: Motility assay on 50 mM β-alanine betaine with wild-type N2 *C. elegans* (Orange), *acr-23* (red), *acr-20* (blue), *deg-3/des-2* (green) and *lgc-41* (pink). n = 30 worms for each genotype, 3 replicates. All values represented as means ± SEM. B: Photograph of N2 after 2 hours on 50 mM β-alanine betaine. Note the lack of sinusoidal wave. C: Photograph of *acr-23* after 2 hours on 50 mM β-alanine betaine. Note the presence of body bends in the worm.

We then treated other *C. elegans* null mutants of the betaine receptors on our β-alanine betaine plates. Fig. 7A shows the effects of 50 mM β*-*alanine betaine on *acr-20 (ok1849), deg-3 (u662)/des-2 (u695)* double mutants and *lgc-41(sy1495)* null mutants. Over the 2-hour period, β-alanine betaine affected the nulls differently: the most resistant worms of these nulls were the *acr-20* null mutants, Fig. 7A: blue line. Only 25% of the worms of the *acr-20* nulls were paralyzed after 2 hours, a result like the *acr-23* null mutants. Both the *acr-20* nulls and *acr-23* nulls were both still capable of body bends. The DEG-3/DES-2 *(u662/u695)* double mutant was less resistant to β*-*alanine betaine compared to *acr-23* and *acr-20*, with 35% of worms being paralyzed after two hours Fig. 7A: green line. The *lgc-41* mutants were even less resistant with 45% of worms being paralyzed Fig. 7A: pink line, after 2 hours. Our results suggest that β-alanine betaine could act on the ACR-23 and ACR-20 receptors but also less potently on the DEG-3/DES-2 heteromeric receptors and LGC-41 receptors.

## DISCUSSION

We detected β-alanine betaine, but not betaine, to be present in the pseudocoelomic fluid of *Ascaris suum* as well as message for members of the DEG-3/DES-2 nAChR subfamily in the body wall and intestine. Injection of β-alanine betaine produced a characteristic ‘pretzel’ coiling paralysis. Application of β-alanine betaine to the *Ascaris* intestine produced a clear calcium signal and depolarized the body wall muscle. Finally, *C. elegans acr-20, acr-23, deg-3/des-2* and *lgc-41* mutants were found to resist the paralytic effects of β-alanine betaine. Together these observations suggest that β-alanine betaine can serve as an endogenous ligand for some members of the DEG-3/DES-2 nAChR subfamily in *A. suum*.

### Betaine in aerobic and β-alanine betaine in anaerobic environments

Betaine may be produced by oxidation of choline or be taken up from the environment: in *C. elegans* choline is first oxidized to betaine aldehyde by choline dehydrogenase and then to betaine, by betaine aldehyde dehydrogenase [6]. Betaine can also be taken up from the environment by a high-affinity transporter, SNF-3 [6]. In parasitic nematodes, betaine has been found to be present in *H. contortus* [15].

We found β-alanine betaine but no detectable betaine in adult *A. suum* which live in a low oxygen environment in the host pig intestine. In hypoxic plants [22], β-alanine betaine is synthesized from β-alanine via methyltransferases: β-alanine is methylated to N-methyl β-alanine, then methylated again to N-N-dimethyl β-alanine and then again to β-alanine betaine. This pathway relies exclusively on methyltransferases and is independent of oxygen, in contrast to choline oxidation pathways required for betaine synthesis. β-alanine betaine is preferentially accumulated in organisms adapted to high saline and hypoxic or anaerobic conditions [23]. Thus, it is possible that *A. suum* in its anaerobic environment produces β*-*alanine betaine rather than betaine as an endogenous ligand.

### Ion-channels activated by betaine and choline in C. elegans and H. contortus

Choline and betaine have been reported in metabolomic studies of *C. elegans* [18, 24]. Both betaine and choline serve different functions, including acting as osmolytes, and in the case of choline and betaine, as endogenous ligands of nAChRs. Betaine and choline selectively activate nematode members of the DEG-3 subfamily of nAChR ion-channels. Expressed *C. elegans* ACR-23 homomeric nAChR channels are activated by betaine (EC_50_ =1.4 mM) but not choline or acetylcholine [6]. *C. elegans* ACR-20 homomeric nAChR channels are selectively activated by betaine (EC_50_ ∼25 µM), less by choline (EC_50_ ∼1.2 mM) and hardly at all by acetylcholine [7, 8]. *C. elegans* DEG-3/DES-2 channels themselves are activated by betaine (EC_50_ ∼ 0.6 mM) and less so by choline (EC_50_ ∼1.8 mM) [8]. Thus, betaine is more potent than choline and acetylcholine on these *C. elegans* channels. Β-alanine betaine has not yet been tested on expressed nAChR channels.

Betaine is also found in the metabolome of *H. contortus* [15]. Studies of *H. contortus* expressed MPTL-1 channels (formally referred to as Hc-ARC-23-H) found that betaine (EC_50_ = 41 µM) activates these channels, and that choline (EC_50_ 1.3 mM) is less potent [7]. Ruferner et al. [25] studied effects of acetylcholine and choline on expressed *H. contortus* DEG-3/DES-2 channels and found that choline had an EC_50_ of 10 mM, but that acetylcholine was a poor agonist even at 10 mM; unfortunately, the effects of betaine on this nAChR were not studied.

Although we found signals for acetylcholine, choline, and β-alanine betaine present in peri-enteric fluid of *Ascaris suum,* we did not find evidence of the presence of betaine. Betaine is produced by oxidation of choline and requires an aerobic environment; the absence of betaine and the presence of β-alanine betaine in *A. suum* may be explained by its anaerobic environment: β-alanine betaine being produced anaerobically [23, 22]). Thus, if betaine functions as both an osmolyte and an endogenous ligand in aerobic nematodes, β-alanine betaine may fulfill analogous roles in anaerobic species such as *A. suum*.

### Effects of monepantel illustrate the therapeutic significance of the DEG-3/DES-2 subfamily

Monepantel (an ADD: amino-acetonitrile derivative) is a class of anthelmintic with activities against nematode parasites of ruminants including *H. contortus, T. colubriformis, Nematodirus spat Higer, Cooperia oncophora* and *Teladorsagia circumcincta.* Monepantel also inhibits movement of *C. elegans* causing hypercontraction of the body wall and anterior pharynx. This hypercontraction is mediated in part by *acr-23* nAChRs because *C. elegans* null-mutants are resistant to ADDs [5]. Furthermore, monepantel has been identified to function as a type 2 positive allosteric modulator (PAM) on members of the DEG-3/DES-2 subfamily [6, 7]. Combinations of monepantel and betaine positively modulate signals in *C. elegans* via ACR-23, ACR-20 and in the *H. contortus* channel MPTL-1 [6, 7], but has no PAM effect on betaine signaling on the DEG-3/DES-2 channels [8]. The action of monepantel is different in *A. suum* where it is not effective therapeutically and behaves as a non-competitive antagonist of ACR-16s and on the levamisole and pyrantel nAChRs [26]. Even high concentrations (10 µM) of monepantel does not produce contraction of *A. suum* body flaps.

### β-alanine betaine as an endogenous ligand in A. suum

Betaine activates members of the DEG-3/DES-2 subfamily in aerobic nematodes. In the absence of detected betaine, we suggest that β-alanine betaine may substitute functionally for betaine in *A. suum* to activate members of the DEG-3/DES-2 subfamily. This view is supported by: 1) the absence of betaine and presence of β-alanine betaine in its peri-enteric fluid in *A. suum*; 2) the absence of an effect of betaine and the pretzel effects of β-alanine betaine following injection in *A. suum*; 3) the increase in cytoplasmic Ca^2+^ in enterocytes following application of β-alanine betaine; 4) the depolarizing effects of β-alanine betaine on *A. suum* somatic muscle cells; and 5) the presence of *acr-20, acr-23, deg-3, des-2,* and *lgc41* in the body wall and intestine enterocytes. We point out however that the aerobic *C. elegans* null mutants of *acr-20 (ok1849), acr-23 (ok 2804), deg-3 (u662)/des-2 (u695)* and *lgc-41(sy1494)* were resistant to β-alanine betaine which suggests that the DEG-3/DES-2 subfamily in *C. elegans* is also sensitive to β-alanine betaine.

## Conclusion

Nematodes have multiple subfamilies of nAChRs. We have identified β-alanine betaine in addition to acetylcholine and choline and suggest that it is a novel ligand in *A. suum*. We found evidence that suggests that β-alanine betaine can act on the DEG-3/DES-2 subfamily of nAChRs. Multiple subfamilies of nAChRs and multiple endogenous cholinergic ligands permit an increase in the complexity of the control the systems that include movement. Given that the DEG-3/DES-2 subfamily of nAChRs is not present in the parasite hosts, it also provides opportunities for the development of selective drugs as anthelmintics.

## Acknowledgement

The authors acknowledge financial support from the National Institute of Allergy and Infectious Diseases (R01AI047194 and R01AI155413) and the E. A. Benbrook Foundation, awarded to RM.

## REFERENCES

1. Jones AK, Davis P, Hodgkin J, Sattelle DB. The nicotinic acetylcholine receptor gene family of the nematode Caenorhabditis elegans: an update on nomenclature. Invert Neurosci. 2007;7(2):129–31. Epub 20070515. doi: 10.1007/s10158-007-0049-z. PubMed PMID: 17503100; PubMed Central PMCID: PMCPMC2972647.

2. Wolstenholme AJ, Neveu C. The interactions of anthelmintic drugs with nicotinic receptors in parasitic nematodes. Emerg Top Life Sci. 2017;1(6):667–73. doi: 10.1042/ETLS20170096. PubMed PMID: 33525839.

3. Buxton SK, Charvet CL, Neveu C, Cabaret J, Cortet J, Peineau N, et al. Investigation of acetylcholine receptor diversity in a nematode parasite leads to characterization of tribendimidine- and derquantel-sensitive nAChRs. PLoS Pathog. 2014;10(1):e1003870. doi: 10.1371/journal.ppat.1003870. PubMed PMID: 24497826; PubMed Central PMCID: PMCPMC3907359.

4. Holden-Dye L, Joyner M, O’Connor V, Walker RJ. Nicotinic acetylcholine receptors: a comparison of the nAChRs of Caenorhabditis elegans and parasitic nematodes. Parasitol Int. 2013;62(6):606–15. Epub 20130315. doi: 10.1016/j.parint.2013.03.004. PubMed PMID: 23500392.

5. Kaminsky R, Ducray P, Jung M, Clover R, Rufener L, Bouvier J, et al. A new class of anthelmintics effective against drug-resistant nematodes. Nature. 2008;452(7184):176–80. PubMed PMID: 992.

6. Peden AS, Mac P, Fei YJ, Castro C, Jiang G, Murfitt KJ, et al. Betaine acts on a ligand-gated ion channel in the nervous system of the nematode C. elegans. Nat Neurosci. 2013;16(12):1794–801. Epub 20131110. doi: 10.1038/nn.3575. PubMed PMID: 24212673; PubMed Central PMCID: PMCPMC3955162.

7. Baur R, Beech R, Sigel E, Rufener L. Monepantel irreversibly binds to and opens Haemonchus contortus MPTL-1 and Caenorhabditis elegans ACR-20 receptors. Mol Pharmacol. 2015;87(1):96–102. Epub 20141028. doi: 10.1124/mol.114.095653. PubMed PMID: 25352042.

8. Hansen TVA, Sager H, Toutain CE, Courtot E, Neveu C, Charvet CL. The *Caenorhabditis elegans* DEG-3/DES-2 Channel is a Betaine-Gated Receptor Insensitive to Monepantel. Molecules. 2022;27(1). Epub 20220105. doi: 10.3390/molecules27010312. PubMed PMID: 35011544; PubMed Central PMCID: PMC8747062.

9. Martin RJ. The gamma-aminobutyric acid receptor of Ascaris as a target for anthelmintics. BiochemSocTrans. 1987;15(1):61–5. PubMed PMID: 40.

10. Williams PDE, Kashyap SS, Robertson AP, Martin RJ. Diethylcarbamazine elicits Ca^2+^ through TRP-2 channels that are potentiated by emodepside in *Brugia malayi* muscles. Antimicrob Agents Chemother. 2023;67(10):e0041923. Epub 20230920. doi: 10.1128/aac.00419-23. PubMed PMID: 37728916; PubMed Central PMCID: PMCPMC10583680.

11. Vasudevamurthy MK, Lever M, George PM, Morison KR. Betaine structure and the presence of hydroxyl groups alters the effects on DNA melting temperatures. Biopolymers. 2009;91(1):85–94. doi: 10.1002/bip.21085. PubMed PMID: 18781629.

12. Brendza KM, Haakenson W, Cahoon RE, Hicks LM, Palavalli LH, Chiapelli BJ, et al. Phosphoethanolamine N-methyltransferase (PMT-1) catalyses the first reaction of a new pathway for phosphocholine biosynthesis in Caenorhabditis elegans. Biochem J. 2007;404(3):439–48. doi: 10.1042/BJ20061815. PubMed PMID: 17313371; PubMed Central PMCID: PMCPMC1896273.

13. Wangchuk P, Yeshi K, Loukas A. Metabolomics and lipidomics studies of parasitic helminths: molecular diversity and identification levels achieved by using different characterisation tools. Metabolomics. 2023;19(7):63. Epub 20230625. doi: 10.1007/s11306-023-02019-5. PubMed PMID: 37356029; PubMed Central PMCID: PMCPMC10290966.

14. Lan W, Xiao X, Nian J, Wang Z, Zhang X, Wu Y, et al. Senolytics Enhance the Longevity of Caenorhabditis elegans by Altering Betaine Metabolism. J Gerontol A Biol Sci Med Sci. 2024;79(11). doi: 10.1093/gerona/glae221. PubMed PMID: 39434620.

15. Pelantová H, Šadibolová M, Žofka M, Matoušková P, Luzarowski M, Krátký J, et al. Metabolomic and proteomic differences in susceptible and benzimidazole-resistant adult females and males of Haemonchus contortus. Vet Res. 2025;57(1):17. Epub 20251224. doi: 10.1186/s13567-025-01698-3. PubMed PMID: 41437124; PubMed Central PMCID: PMCPMC12849590.

16. Yeshi K, Creek DJ, Anderson D, Ritmejerytė E, Becker L, Loukas A, et al. Metabolomes and Lipidomes of the Infective Stages of the Gastrointestinal nematodes,. Metabolites. 2020;10(11). Epub 20201106. doi: 10.3390/metabo10110446. PubMed PMID: 33171998; PubMed Central PMCID: PMCPMC7694664.

17. Martin RJ, Robertson AP, Buxton SK, Beech RN, Charvet CL, Neveu C. Levamisole receptors: a second awakening. Trends Parasitol. 2012;28(7):289–96. doi: 10.1016/j.pt.2012.04.003. PubMed PMID: 22607692; PubMed Central PMCID: PMCPMC3378725.

18. Liu F, Li T, Gong H, Tian F, Bai Y, Wang H, et al. Structural insights into the molecular effects of the anthelmintics monepantel and betaine on the Caenorhabditis elegans acetylcholine receptor ACR-23. EMBO J. 2024;43(17):3787–806. Epub 20240715. doi: 10.1038/s44318-024-00165-7. PubMed PMID: 39009676; PubMed Central PMCID: PMCPMC11377560.

19. Hardege I, Morud J, Yu J, Wilson TS, Schroeder FC, Schafer WR. Neuronally produced betaine acts via a ligand-gated ion channel to control behavioral states. Proc Natl Acad Sci U S A. 2022;119(48):e2201783119. Epub 20221121. doi: 10.1073/pnas.2201783119. PubMed PMID: 36413500; PubMed Central PMCID: PMCPMC9860315.

20. Xiong H, Pears C, Woollard A. An enhanced C. elegans based platform for toxicity assessment. Sci Rep. 2017;7(1):9839. Epub 20170829. doi: 10.1038/s41598-017-10454-3. PubMed PMID: 28852193; PubMed Central PMCID: PMCPMC5575006.

21. O’Reilly LP, Luke CJ, Perlmutter DH, Silverman GA, Pak SC. C. elegans in high-throughput drug discovery. Adv Drug Deliv Rev. 2014;69-70:247–53. Epub 20131212. doi: 10.1016/j.addr.2013.12.001. PubMed PMID: 24333896; PubMed Central PMCID: PMCPMC4019719.

22. Parthasarathy A, Savka MA, Hudson AO. The Synthesis and Role of β-Alanine in Plants. Front Plant Sci. 2019;10:921. Epub 20190718. doi: 10.3389/fpls.2019.00921. PubMed PMID: 31379903; PubMed Central PMCID: PMCPMC6657504.

23. Hanson AD, Rathinasabapathi B, Chamberlin B, Gage DA. Comparative Physiological Evidence that beta-Alanine Betaine and Choline-O-Sulfate Act as Compatible Osmolytes in Halophytic Limonium Species. Plant Physiol. 1991;97(3):1199–205. doi: 10.1104/pp.97.3.1199. PubMed PMID: 16668509; PubMed Central PMCID: PMCPMC1081142.

24. Wan QL, Shi X, Liu J, Ding AJ, Pu YZ, Li Z, et al. Metabolomic signature associated with reproduction-regulated aging in. Aging (Albany NY). 2017;9(2):447–74. doi: 10.18632/aging.101170. PubMed PMID: 28177875; PubMed Central PMCID: PMCPMC5361674.

25. Rufener L, Baur R, Kaminsky R, Maser P, Sigel E. Monepantel allosterically activates DEG-3/DES-2 channels of the gastrointestinal nematode Haemonchus contortus. Mol Pharmacol. 2010;78(5):895–902. Epub 2010/08/04. doi: mol.110.066498 [pii] 10.1124/mol.110.066498. PubMed PMID: 20679419.

26. Abongwa M, Marjanovic DS, Tipton JG, Zheng F, Martin RJ, Trailovic SM, et al. Monepantel is a non-competitive antagonist of nicotinic acetylcholine receptors from Ascaris suum and Oesophagostomum dentatum. Int J Parasitol Drugs Drug Resist. 2018;8(1):36–42. Epub 20171216. doi: 10.1016/j.ijpddr.2017.12.001. PubMed PMID: 29366967; PubMed Central PMCID: PMCPMC5963102.

